# The genome of Przewalski’s horse (*Equus ferus przewalskii*)

**DOI:** 10.1101/2024.02.20.581252

**Authors:** Nicole Flack, Lauren Hughes, Jacob Cassens, Maya Enriquez, Samrawit Gebeyehu, Mohammed Alshagawi, Jason Hatfield, Anna Kauffman, Baylor Brown, Caitlin Klaeui, Islam F. Mabrouk, Carrie Walls, Taylor Yeater, Anne Rivas, Christopher Faulk

## Abstract

The Przewalski’s horse (*Equus ferus przewalskii*) is an endangered equid native to the steppes of central Asia. After becoming extinct in the wild, multiple conservation efforts convened to preserve the species including captive breeding programs, reintroduction and monitoring systems, protected lands, and cloning. Availability of a highly contiguous reference genome is essential to support these continued efforts. We used Oxford Nanopore sequencing to produce a scaffold-level 2.5 Gb nuclear assembly and 16,002 bp mitogenome from a captive Przewalski’s mare. All assembly drafts were generated from 111 Gb of sequence from a single PromethION R10.4.1 flow cell. The mitogenome contained 37 genes in the standard mammalian configuration and was 99.63% identical to the domestic horse (*Equus caballus*). The nuclear assembly, EquPr2, contained 2,146 scaffolds with an N50 of 85.1 Mb, 43X mean depth, and BUSCO quality score of 98.92%. EquPr2 successfully improves upon the existing Przewalski’s horse reference genome (Burgud), with 25-fold fewer scaffolds, a 166-fold larger N50, and phased pseudohaplotypes. Modified basecalls revealed 79.5% DNA methylation and 2.1% hydroxymethylation globally. Allele-specific methylation analysis between pseudohaplotypes revealed 226 differentially methylated regions (DMRs) in known imprinted genes and loci not previously reported as imprinted. The heterozygosity rate of 0.165% matches previous estimates for the species and compares favorably to other endangered animals. This improved Przewalski’s horse assembly will serve as a valuable resource for conservation efforts and comparative genomics investigations.

## Introduction

The Przewalski’s horse (*Equus ferus przewalskii*), also called the tahki, is an endangered equid native to central Asia [1] whose lineage diverged from the domestic horse (*Equus caballus*) tens of thousands of years ago [2, 3, 4, 3, 5]. *E. f. przewalskii* has a distinct short and stocky build with dun coloring and an erect mane. These horses were initially native to the steppes of central Asia, and by the 19th century inhabited only Mongolia, Tibet, and China [6, 7]. Introgression from the domestic horse, loss of natural habitat and resources, harsh climates, and hunting contributed to a severe population bottleneck and subsequent extinction in the wild in the 1960s [7, 8]. All current Przewalski’s horses are descendants of 12 wild-caught individuals and several domesticated horses [6, 7, 9]. Massive conservation efforts have focused on preserving the genetic diversity of this endangered species [10, 11, 12, 13, 14, 15].

Targeted captive breeding and management programs, including published studbooks since 1959, have increased the Przewalski’s horse population to over 2,000 individuals [14, 9, 3]. Reintroduction efforts starting in the 1980s have successfully established wild herds in protected lands of Mongolia, China and Kazakhstan [16, 9, 15, 17, 18, 19, 20, 21], and have improved the species’ status from critically endangered to endangered [1]. Recent cloning efforts through the Przewalski’s Revive & Restore Project have produced two males from a cryopreserved cell line in the San Diego Zoo Wildlife Alliance Frozen Zoo [22, 23, 24], further bolstering captive breeding efforts.

Przewalski’s horses possess additional chromosomes (2n=66) when compared to the domestic horse (2n=64) [25, 26, 27, 28]; this chromosome difference is thought to be derived from a Robertsonian translocation event [29, 30]. Przewalski’s and domestic horse crosses produce viable, fertile offspring with odd numbered chromosomes (n=65), in contrast to the infertile offspring of domestic horse and donkey (*Equus asinus*) crosses [31]. Investigation of the differences between domestic and Przewalski’s horse chromosome structures may facilitate improved understanding of chromosome fusion.

Recent technological advances have increased accessibility and reduced costs for generating high-quality whole genome sequencing data [32]. Long-read Oxford Nanopore sequencing provides genomic and epigenomic data simultaneously without amplification or bisulfite conversion [33]. Two major limitations of the technology, high per-base error rates and homopolymer indels, have been addressed with improved basecalling models, self-correction algorithms, and adequate depth of coverage [34]. In addition to direct capture of epigenetic base modifications, pseudohaplotype phasing permits heterozygosity estimation and allele-specific DNA methylation analysis; strict allele-specific methylation is a signature of genomic imprinting, where genes are monoallelically expressed based on parent of origin [35, 36]. The epigenome is also relevant to species evolutionary biology due to the increased mutation rate of methylated cytosines via deamination [37, 38, 39, 40, 41, 42].

Here, we highlight the exclusive use of Oxford Nanopore sequencing reads to provide a high-quality, highly contiguous diploid nuclear genome assembly, updated mitogenome, and DNA methylation analysis for the endangered Przewalski’s horse.

## Methods

### Sample collection

Varuschka, a 10-year-old captive-bred Przewalski’s mare (Figure 1), was subject to routine veterinary care under anesthesia during which 10 ml of whole blood was collected by zoo veterinarians. The Minnesota Zoo has been active in Przewalski’s horse breeding and management, with over 50 foals born since the 1970s, and contributed a stallion to reintroduction efforts in Mongolia’s Hustai National Park [43, 44].

**Figure 1.**
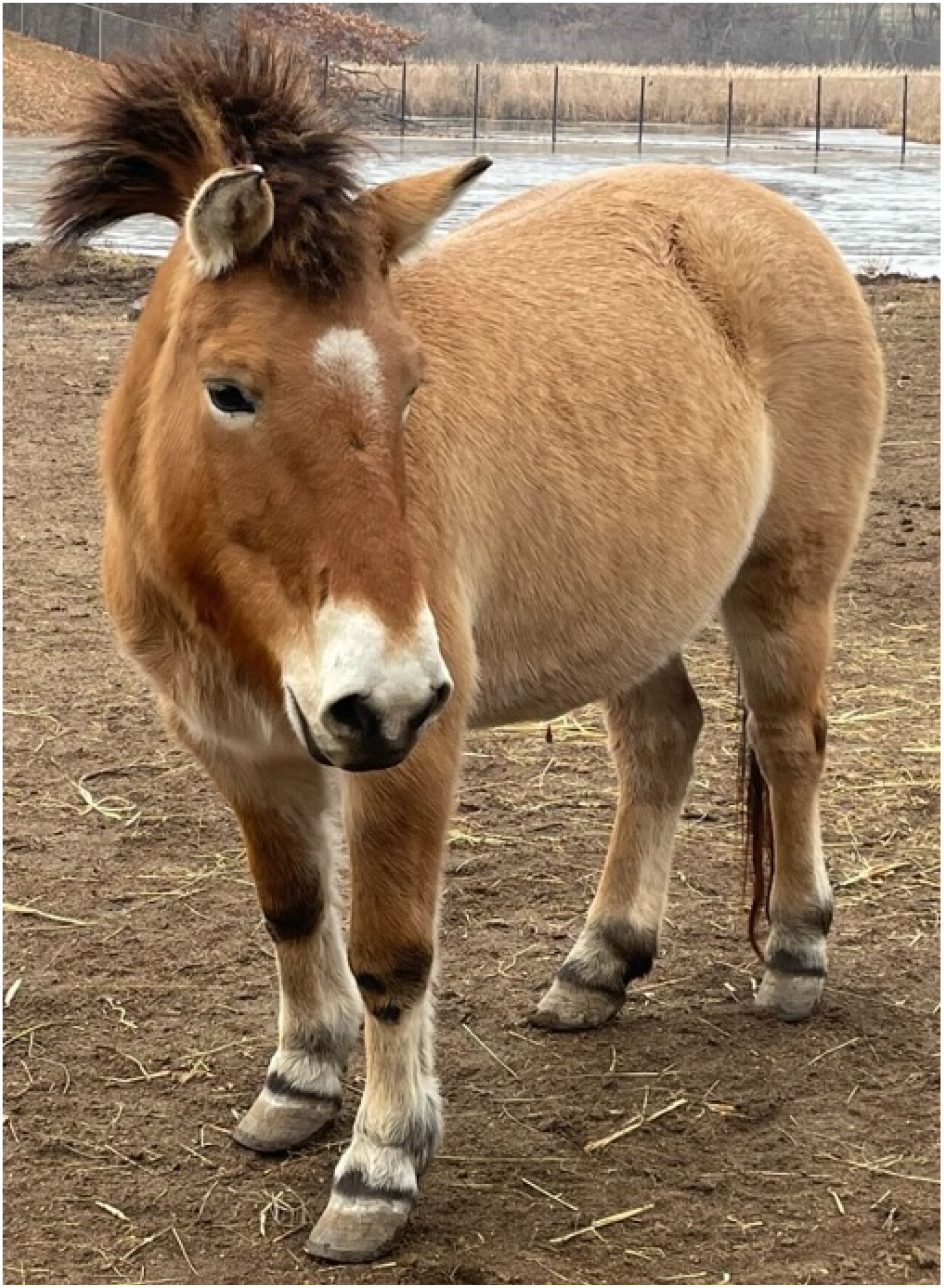
*Equus ferus przewalskii* specimen photo depicting Varuschka, the 10-year-old captive-bred mare sampled for genome assembly. Image courtesy of the Minnesota Zoo.

### DNA extraction and sequencing

Genomic DNA was extracted from blood using a MagAttract Blood DNA/RNA Kit (Qiagen, Venlo, Netherlands) according to manufacturer’s instructions. Sequencing was performed on a P2 Solo instrument (Oxford Nanopore Technologies, Oxford, UK) using a single PromethION R10.4.1 flow cell. We created two libraries using the LSK-114 ligation sequencing kit for native DNA sequencing. For the first library, 3 μg of DNA in 100 μl elution buffer was sheared by passage through a 28 gauge needle 30 times, library prepped, then split into three aliquots of 15 μl each. For the second library, unsheared DNA was eluted and prepped as a single 15 μl library. The first aliquot of the sheared library was loaded onto the flowcell and run for 24 hours, after which the flowcell was washed using the manufacturer’s wash kit. The remaining two sheared aliquots were loaded and sequenced in the same fashion on days 2 and 3, respectively. After sequencing the three sheared library aliquots, the unsheared library was loaded into the flowcell and run for 24 hours. Data were collected using 5 kHz minKNOW version 23.07.12 (Oxford Nanopore Technologies, Oxford, UK).

Raw nanopore data from both libraries were basecalled together using Dorado v0.4.3 (https://github.com/nanoporetech/dorado) with “super accuracy” model dna_r10.4.1_e8.2_400bps_sup@v4.2.0; base modifications called simultaneously with flag `--modified-bases 5mC_5hmC`. Read quality was assessed using the Nanoq package (https://github.com/esteinig/nanoq).

### Genome assembly

Detailed computational methods are available in Supplementary File 2. The genome was *de novo* assembled using Flye v2.9 [45], followed by polishing using Medaka v1.6.0 (https://github.com/nanoporetech/medaka). Duplicate contigs were removed using Purge Dups v1.2.6 (https://github.com/dfguan/purge_dups). Additional manual curation was performed to remove contigs with less than 15X or greater than 500X mean coverage. One contig representing the mitochondrial genome was also removed at this stage. The resulting draft assembly was scaffolded onto the reference domestic horse genome, EquCab3.0 (GCF_002863925.1), using the RagTag package v2.1.0 [46]. Gaps were closed using TGS-GapCloser v1.2.1 [47].

Scaffolds for *E. ferus przewalskii* were assigned chromosome names based on homology to *E. caballus* synteny from EquCab3.0 as demonstrated previously [30]. The homologous *E. f. przewalskii* scaffold to *E. caballus* chromosome 5 was split on its scaffold gaps with AWK, and the largest blocks syntenic to the p- and q-arms were labeled with their chromosome names (Supplementary File 1). The remaining smaller contigs syntenic to EquCab3.0 chromosome 5 were labeled as ChrUns in the scaffolded assembly. The FCS-adapter tool from the NCBI Foreign Contamination Screening program suite was used to detect and remove adapter and vector contamination from the final haplotype assemblies (https://github.com/ncbi/fcs).

The quality of all draft assemblies was evaluated by detecting Benchmarking Universal Single-Copy Ortholog (BUSCO) genes within the *Cetartiodactyla* lineage [48]. We used the Compleasm package to calculate BUSCO scores as it is a faster, more accurate implementation of BUSCO [49]. We combined Compleasm’s single and duplicate BUSCO counts to provide a direct comparison to the standard BUSCO program’s ‘complete’ value. On average, Compleasm detected 3% more BUSCOs than the BUSCO tool v3.4.7 (Supplementary File 1). We calculated assembly N50, L50, and other statistics using the packages Assembly-Stats (https://github.com/sanger-pathogens/assembly-stats) and the Quality Assessment Tool for Genome Assemblies (QUAST v5.2.0) [50], using EquCab3.0 as reference for the latter. Assembly statistics were compared to other publicly available equid genomes including the preexisting Przewalski’s horse (Burgud), domestic horse (EquCab3.0), and plains zebra genome (*Equus quagga*; UCLA_HA_Equagga_1.0) references.

### Methylation

Global DNA methylation and hydroxymethylation (5mC and 5hmC) at cytosine-guanine dinucleotides (CpGs) was determined using the modified base information stored in the basecalled BAM files, i.e., modBAMs. Since the original basecalling was performed prior to a reference existing, Dorado stored modified bases per-read as unmapped modBAMs. These were concatenated together and aligned to the final scaffold-level assembly using Minimap2 [51, 52]. The resulting mapped modBAMs were converted to bedMethyl format using Modkit v0.2.2 (https://github.com/nanoporetech/modkit) with global 5mC and 5hmC percentages summarized using AWK (https://www.gnu.org/software/gawk/manual/gawk.html). For allele-specific DNA methylation, Modkit was applied to the phased modBAM generated by variant calling to generate two bedMethyl files separated by haplotag. Count filtering, normalization, tiling, and differential methylation testing were performed with MethylKit [53, 54]. Nearest genes to significant regions were found with Bedtools [55]. Differentially methylated regions (DMRs) between haplotypes were visualized with Methylartist [56].

### Repeats

RepeatMasker v4.1.4 (https://www.repeatmasker.org/) was used to identify repetitive sequences with the complete Dfam library v3.6 (https://www.dfam.org/home) as described previously [57, 58]. For consistency of comparison, Repeatmasker was also run locally on the existing equid reference genomes with the same parameters.

### Variant calling and diploidization

Clair3 v1.05 [59] was run to determine the number and type of variants. Heterozygosity was calculated by counting the number of variants divided by the total genome size. Variants were phased with Whatshap [60] and haplotagged with Longphase [61]. BCFTools was used to swap out the phased variants in the primary assembly to generate the secondary pseudohaplotype assembly (https://github.com/samtools/bcftools).

### Gene annotation

Homology-based gene prediction was performed with Gene Model Mapper (GeMoMa, https://doi.org/10.1007/978-1-4939-9173-0_9) using EquCab3.0 transcripts as the reference. BUSCO was used in protein mode to assess gene prediction accuracy and completeness.

### Mitochondrial assembly

The mitogenome was extracted from the *E. ferus przewalskii* assembly using MitoHiFi v3.2 [62, 63]. The program identifies mitogenome contigs by comparison to known mitogenomes from related species; in this case we used the EquCab3.0 mitogenome (NC_001640.1). MitoHiFi also circularizes and annotates the putative mitogenome contig.

## Results and Discussion

### Assembly

We generated a total of 111 Gb of DNA sequencing data for Przewalski’s horse with a read N50 of 10,829 bp and mean quality of Q18.49. The quality of each draft assembly was assessed with parameters including N50 (i.e., length of the shortest contig at 50% of the total assembly length), L50 (i.e., smallest number of contigs whose length sum to 50% of the total assembly length), and the count of benchmark universal single-copy ortholog (BUSCO) genes. Our initial Flye run yielded a 2.59 Gb draft assembly with 6,808 contigs and an N50 of 13.6 Mb (Table 1). With an L50 of 55, the majority of the genome was assembled into relatively few contigs representing large portions of chromosomes.

**Table 1.**
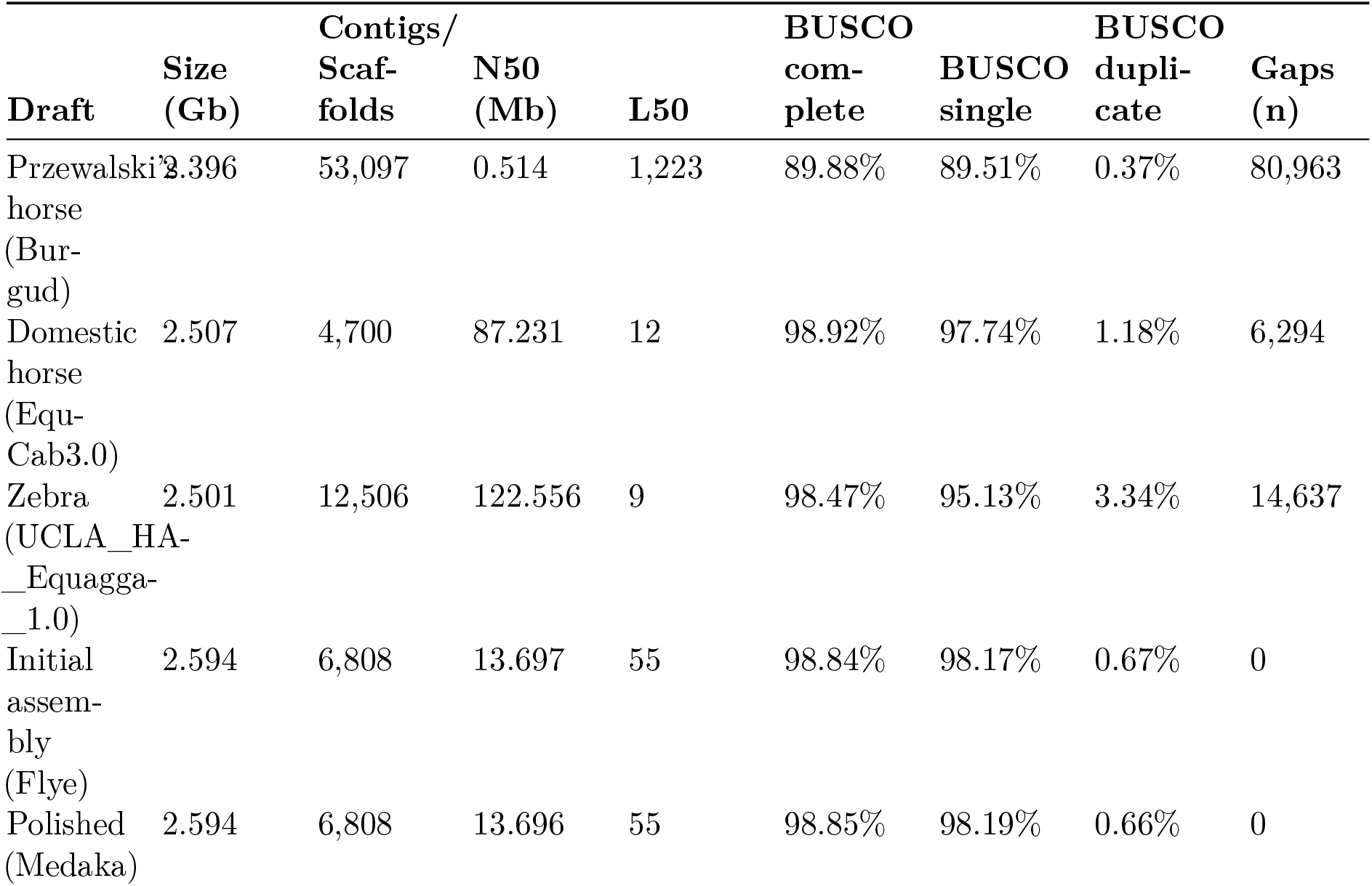
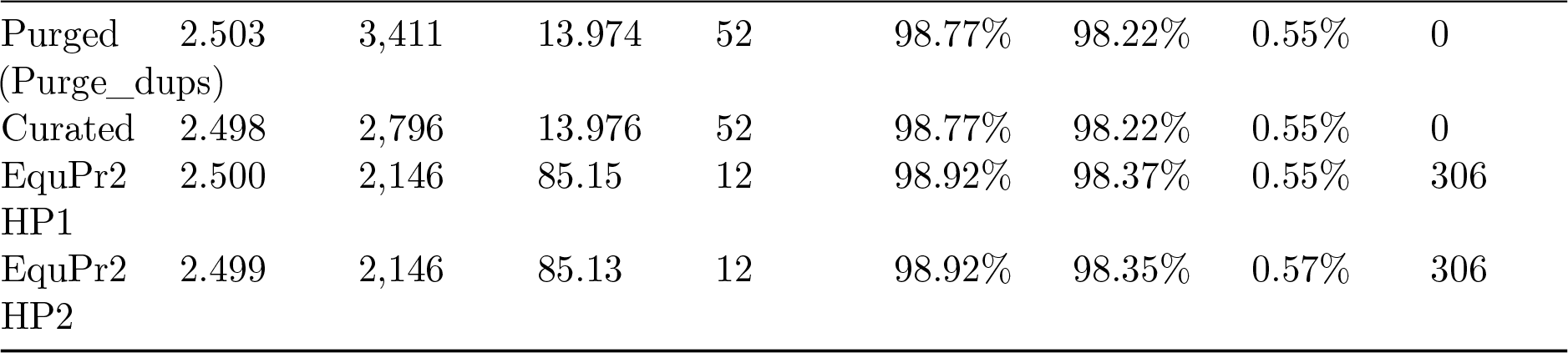
Contiguity and quality statistics for draft *E. ferus przewalskii* assemblies, final EquPr2 assemblies, and existing equid reference genomes. Benchmark universal single-copy ortholog (BUSCO) scores were calculated with the Compleasm tool [49]. HP: pseudohaplotype.

The initial assembly was polished, purged of duplicates, and manually curated to remove contigs with coveage below 15X and 500X as they are unlikely to represent single-copy nuclear regions [64]. This procedure resulted in the removal of 4,012 contigs and 96 Mb of sequence. Polishing with Medaka increased the complete BUSCO score by reducing the percentage of duplicate BUSCOs by 0.01%. In our initial draft, the duplicate BUSCO count was already low at less than 1% of the total; haplotig purging reduced the number of duplicate BUSCOs by 0.11% and increased N50 from 13.70 Mb to 13.97 Mb.

We scaffolded the curated assembly onto the domestic horse genome (EquCab3.0); gaps were filled with reads placed by TGS-GapCloser where spanning reads could be identified. This final assembly, EquPr2, was 2.50 Gb in length with a chromosome-level N50 of 85.1 Mb and L50 of 12. EquPr2 contained 306 gaps spanning 2.03 Mb, a 5-fold and 11-fold reduction in gap length compared to EquCab3.0 and Burgud, respectively (Table 1). The contiguity of EquPr2 also matches EquCab3.0’s L50 of 12, in contrast to Burgud’s L50 of 1,223 (Figure 2).

**Figure 2.**
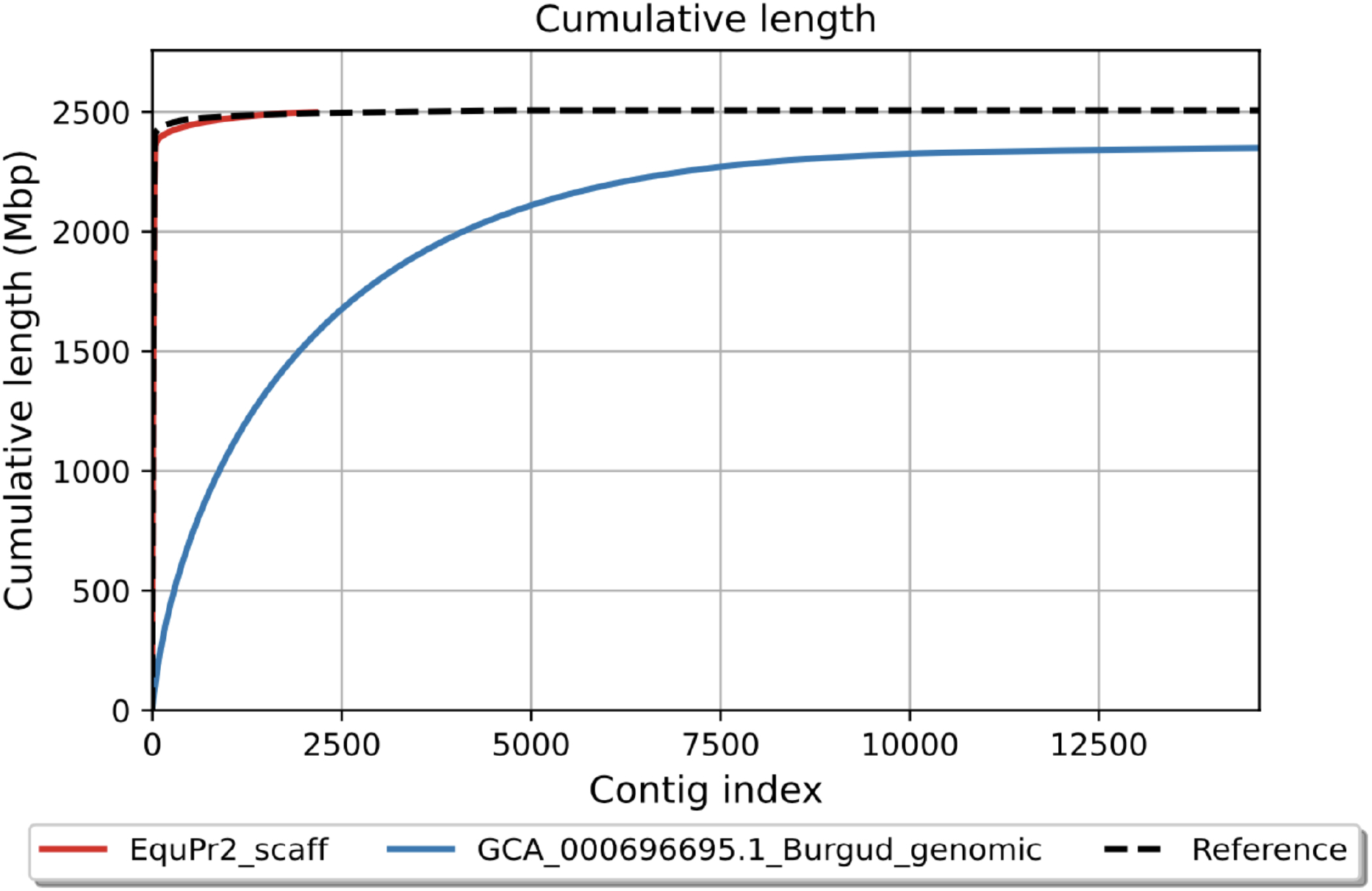
Cumulative length of EquPr2 scaffolds.

Our Przewalski’s horse genome had a BUSCO completeness score of 98.92%, improving from 89.88% in the previous Przewalski’s horse assembly with 25-fold fewer scaffolds and a 166-fold increase in N50. It is important to deconstruct the complete BUSCO score reported by most new genome assemblies into its component parts of single and duplicate copy percentages.

Examining only the completeness score obscures the presence of haplotig missassemblies represented by high duplicate counts. For instance, the reference horse genome, EquCab3.0, is a high-quality, highly contiguous assembly with a complete BUSCO score of 98.92%, the same as our Przewalski’s horse assembly EquPr2. However EquCab3.0 has a duplicate rate of 1.18% versus EquPr2’s duplicate rate of 0.55%. Similarly, the plains zebra genome, UCLA_HA_Equagga1.0, is even more contiguous with an L50 of 9 but has a 3.34% BUSCO duplication rate. While these differences are relatively small, duplication misassemblies have contributed to erroneous gene gain and gene family expansion findings in high-heterozygosity vertebrate genomes [65].

The Przewalski’s horse has 33 sets of chromosomes versus the domestic horse’s 32 due to a Robertsonian translocation where *E. caballus* chromosome 5 is homologous to *E. f. przewalskii* chromosomes 23 and 24 [30]. Other EquPr2 chromosomes were named based on homology to domestic horse consistent with previous karyotyping [66, 67]. We manually split the EquPr2 scaffold homologous to *E. caballus* chromosome 5 at every gap from the contig-level assembly and named the largest contigs as chromosomes 23 and 24 based on homology to the p and q arms of its species ortholog. Due to lack of positional certainty, the remaining contigs mapping to EquCab3.0’s chromosome 5 were named as chromosome unknown (ChrUn).

### Repetitive DNA

We used RepeatMasker to detect repetitive element content within the new assembly and compared it to the previous assembly, Burgud, and the domestic horse (Table 2). Given the genetic similarity of Przewalski’s horse to the domestic horse, global transposon content is nearly identical across all categories. Identification of single repeat insertions facilitated by a more contiguous Przewalski’s horse reference may be valuable for future comparative genomics investigations of these two closely related species.

**Table 2.** Repetitive DNA content for EquPr2, the existing Przewalski’s horse reference (Burgud), and the domestic horse reference (EquCab3.0). Only RepeatMasker categories reaching >1% genomic content are shown; smaller families are collapsed into the relevant parent category.

| RepeatMasker | EquPr2 (%) | Burgud (%) | EquCab3.0 (%) |
| --- | --- | --- | --- |
| **Retroelements** | 30.89 | 31.66 | 32.21 |
| SINEs: | 3.55 | 3.70 | 3.66 |
| LINES: | 21.35 | 21.65 | 22.27 |
| L2/CR1/Rex | 5.26 | 5.47 | 5.43 |
| L1/CIN4 | 15.88 | 15.96 | 16.63 |
| LTRs: | 6.00 | 6.32 | 6.28 |
| Retroviral | 5.57 | 5.88 | 5.84 |
| **DNA transposons** | 3.44 | 3.62 | 3.56 |
| hobo-Activator | 2.6 | 2.74 | 2.69 |
| **Total repeats** | **34.36** | **35.32** | **35.8** |

### Heterozygosity

Given the extreme population bottleneck that occurred during the near-extinction of Przewalski’s horse, it is critical to understand the genetic diversity remaining for captive breeding efforts. To this end and due to lack of samples for Varuschka’s sire and dam, we called and phased EquPr2 variants to build an alternate pseudohaplotype assembly. Variant calling with Clair3 found 4,114,297 variants, the majority of which were single-nucleotide variants (SNVs). Heterozygosity was estimated to be 0.165. This level of genetic diversity is concordant with previous microarray data from nine Przewalski’s horses where average heterozygosity was 0.168, the highest estimate among nine members of *Hippomorpha* excluding the domestic horse [68].

### Gene annotation

We detected 21,552 putative genes in EquPr2 with the *ab initio* gene prediction tool Gene Model Mapper (GeMoMa) [69] using the EquCab3.0 annotation (GCF_002863925.1) as reference. The resulting protein BUSCO score was 85.6% complete (84.6% single copy and 1.0% duplicates); this score appears to be near GeMoMa’s maximum performance based on previous demonstration of similar results for existing high-quality reference genomes [64]. The EquPr2 annotation would likely be significantly improved with by the application of NCBI’s annotation pipeline, which requires the availability of RNA-seq data.

### Mitochondrial genome

The mitochondrial genome was extracted from the assembly and characterized with MitoHiFi, a tool designed for long-read sequencing mitochondrial contig identification and annotation [63]. The resulting mitogenome for EquPr2 was 16,002 bp, 8 bp shorter than EquCab3.0, possibly resulting from incomplete assembly. It contained 37 genes in the standard mammalian configuration and was 99.63% identical to *E. caballus* isolate TN9488 (Supplementary File 1).

### DNA methylation

Oxford Nanopore sequencing can natively detect base modifications including 5-methylcytosine (5mC). This feature has been applied in previous mammalian genome assemblies to evaluate allele-specific DNA methylation and identify known and putative novel imprinted genes [64]. Globally, we found that whole blood leukocyte DNA methylation in Przewalski’s horse was 79.5% methylated and 2.1% hydroxymethylated at CpG sites genome-wide. After filtering to include CpGs with at least 10x coverage in both pseudohaplotypes, there were 18,928,678 sites available to test for allele-specific differential methylation. Counts were tiled into 100 bp windows; windows containing at least 10 CpGs with an absolute difference in methylation >=50% and Benjamini-Hochberg-adjusted p-value < 0.05 were deemed differentially methylated regions (DMRs). With these parameters, we identified 226 DMRs between EquPr2 pseudohaplotyes with a mean absolute methylation difference of 64.1% (Figure 3). Nearest features to DMRs included known imprinted genes (e.g., *IGF2R, INPP5F, PEG3, DIRAS3*) and loci not previously reported as imprinted (e.g., *DLG3*). Seventy-seven DMRs (34.1%) directly overlapped predicted genes. As a consequence of random X-chromosome inactivation (XCI) in a female animal, 111 of the 226 DMRs (49.1%) were located on ChrX. Imprinted genes are unique to mammals and strongly influence growth, making this information valuable for future investigations of Przewalski’s horse evolution, comparative genomics, and conservation.

**Figure 3.**
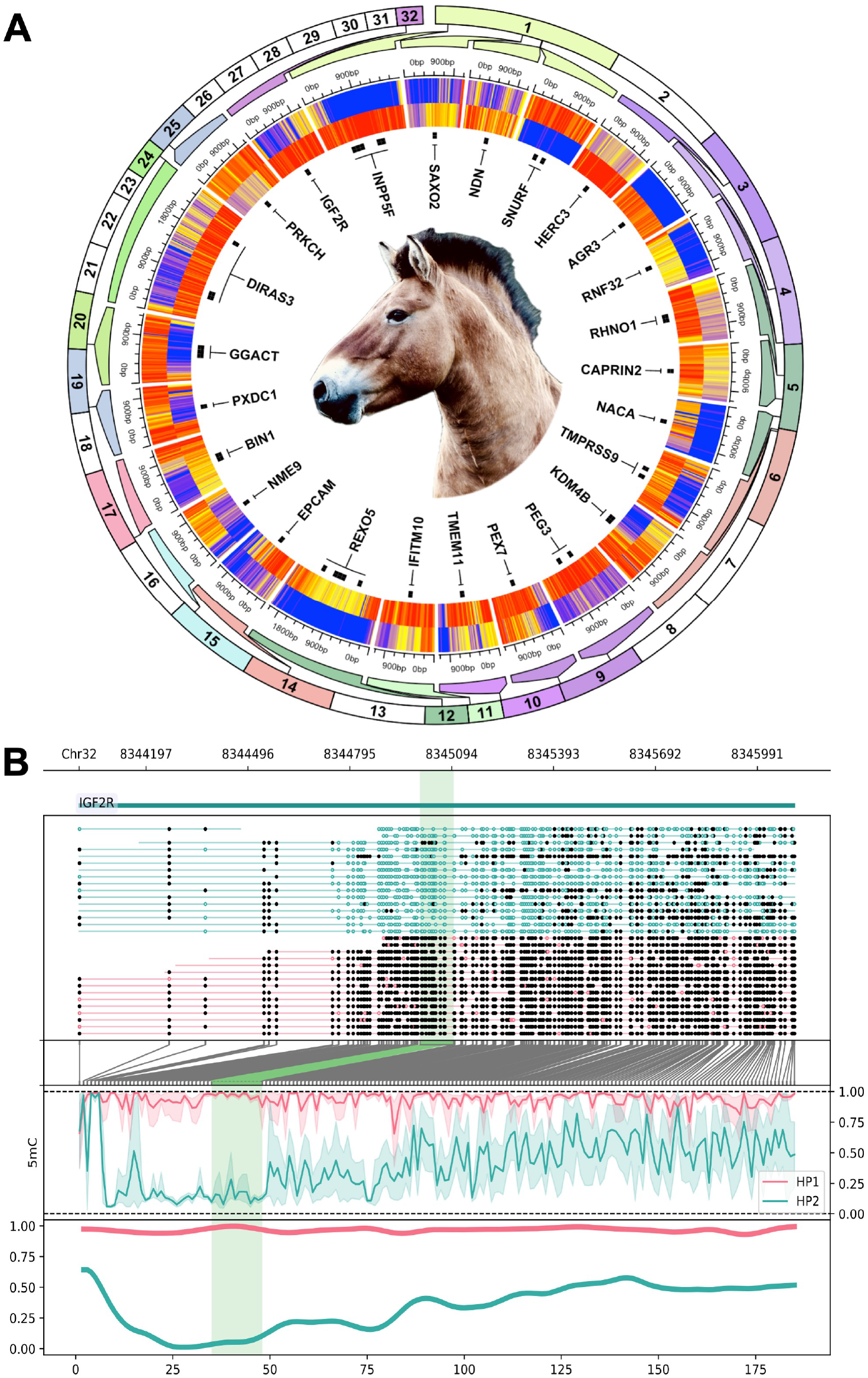
Genome-wide allele-specific DNA methylation analysis for EquPr2. (A) Circular heatmap of 115 differentially methylated regions (DMRs) identified between EquPr2 pseudohaplotypes and labeled by nearest gene symbol. Regions are zoomed from their chromosomal position as indicated by outer track color. Each heatmap row is a pseudohaplotype and each column is a single CpG within the DMR. Methylation values range from 0% (blue) to 100% (red). (B) Example DMR overlapping the known imprinted gene *IGF2R*. Filled circes are methylated CpGs and open circles are unmethylated CpGs.

## Conclusions

The availability of a high-quality reference genome is imperative for improved understanding of the genetic diversity of Przewalski’s horse. The lineage of the horse sampled for this paper’s holotype animal, Varuschka, traces back to founders from Mongolia; her dam was imported from the Cologne Zoo in Germany as a part of the Species Survival Plan (SSP), and her sire was transferred from the Smithsonian Zoo. Here, we provide a 2.5 Gb nanopore-only genome assembly for Przewalski’s horse with an improved BUSCO score of 98.92%, 25-fold fewer scaffolds, and a 166-fold increase in N50 compared to the existing reference genome. Modified basecalls additionally facilitated allele-specific methylation analysis; significantly differentially methylated regions included known imprinted genes and potential novel loci. This genome will aid Przewalski’s horse conservation by providing a higher quality, more contiguous foundation for captive breeding, population genomics, and other efforts.

## Author Contributions

This work was a collaborative effort by the members of graduate course ANSC 8520 taught by Dr. Faulk in the Department of Animal Science at the University of Minnesota in Fall 2023. Students were Cassens, Enriquez, Gebeyehu, Alshagawi, Hatfield, Kauffman, Klaeui, Mabrouk, Walls, Brown, and Dr. Yeater. Students performed analyses and provided text for the manuscript. Drs. Hughes, Martinson, Rivas, and Koch provided the horse sample, analytical guidance, and edited the manuscript. Dr. Flack analyzed methylation data, provided text for the manuscript, and edited the manuscript. Dr. Faulk performed analyses and edited the manuscript. All authors have read and approved the manuscript prior to submission.

## Data availability

Supplementary File 2 contains code used to generate data. The diploid assembly and mitogenome are available under NCBI umbrella BioProject PRJNA1073944 and submissions JAZHEL000000000 and JAZHEM000000000. Variant calls and other supplementary material are available on figshare.

## Acknowledgments

We thank the Minnesota Zoo for generously donating the sample from Varuschka, as well as the Animal Care and Animal Health staff who provided sample collection assistance and knowledge of Przewalski’s horse history. We thank Krishona Martinson for helpful discussions on the biology of the horse.

