## Supplementary File 2 for "The genome of Przewalski’s horse (*Equus ferus przewalskii*)"

### Supplemental File 2

**The genome of Przewalski's horse (*Equus ferus przewalskii*)**

#### Table of contents

**Genome Assembly Pipeline** **2**

**Authors** **2**

**Species and Sample** **3**

**Assembly** **3**

**Scaffolding (ragtag)** **6**

**Scaffolding (ntLink)** **7**

**Gap Closing** **7**

**Global Methylation** **9**

**RepeatMasking** **10**

**Annotation** **10**

**Variant Calling & Phasing** **10**

**DNA Methylation** **11**

**BUSCO** **20**

**Quast** **21**

### Genome Assembly Pipeline

This is the Faulk lab pipeline for *de novo* animal genome assembly.

### Authors

Nicole Flack, Samrawit Gebeyehu, Carrie Walls, Lauren Hughes, Krishona Martin, Mohammed Alshagawi, Anna Kauffman, Baylor Brown, Taylor Yeater, Jacob Cassens, Maya Enriquez, Caitlin Klaeui, Jason Hatfield, Islam F. Mabrouk, Chris Faulk

#### Species and Sample

Sample origin: Varuschka, a captive Przewalksi's mare housed at the Minnesota Zoo. DNA source: Whole blood. Extraction method: Zymo Sequencing instrument: Oxford Nanopore P2 Solo, flowcell R10.4.1 Sequencing Run: 1 flow cells had 3 libraries loaded, each preceded by one wash (wash kit 004). Library prep: Aliquots of 3 ug of DNA from whole blood were each end prepped using NEBNext FFPE Repair Mix (M6630), NEBNext Ultra II End repair/dA-tailing Module (E7546), and NEBNext Quick Ligation Module (E6056) and subsequently library prepped using the ONT supplied SQK- LSK114 kit. The resulting library was eluted in 45 ul of elution buffer and aliquotted into three 15 ul libraries for loading.

#### Assembly

We assembled the genome with the following properties:

- Diploid
- Mitochondrial Genome
- DNA methylation and hydroxymethylation

##### Methods

**Example**: Computational environment and software sequencing computer was built with an AMD Ryzen 3900x processor with 12 cores, 24 threads, 64 Gb RAM and a 4 Tb SSD, running Ubuntu Linux 22.04. For basecalling we added two GPUs, both GeForce RTX 4090. Genome assembly with flye was performed on a 128 core cluster node with 2 Tb of RAM.

##### Basecalling

Dorado v0.4.3 was used to call bases and CpG modifications. The naming convention is that `$FILE.mod.bam` indicates a bam file containing sequencing reads with modification calls and `$FILE.modmapped.bam` contains reads with modifications *mapped* to a genome.

```
# NOTE THAT WE CALLED ALL C SITES, NOT JUST CPG!
dorado basecaller dna_r10.4.1_e8.2_400bps_sup@v4.2.0 Sample_pod5_directory
↪  --modified-bases 5mC_5hmC -r > p-horse-total.bam
```

#### QC on Reads

Quality Control on Reads

```
# Basic read statistics with nanoq (faster than seqkit).
cargo install nanoq
samtools fastq <file> | nanoq -vvv -s
# -or-
nanoq -vvv -s -i file.fastq.gz

# Nanoplot with unmapped bam
NanoPlot -t 12 -o nanoplot-out --ubam file.bam
```

#### Flye Assembly

Flye typically gives the best assemblies. Flye was run alternatively with and without the –asm-coverage flag to limit coverage across the assembly. Assemblies were compared with the full read set and a subset of filtered reads.

##### Install

Flye was installed from github via conda

```
conda env create -n flye -c bioconda flye=2.9.1
conda activate flye
```

##### Slurm

```
#!/bin/bash -l
#SBATCH -A faulkc
#SBATCH --time=24:00:00
#SBATCH -p ag2tb
#SBATCH --ntasks=128
#SBATCH --mem=1995g
#SBATCH --mail-type=ALL
#SBATCH --mail-user=

Flye/bin/flye --nano-hq phorse_varuska_both_uniq.fastq  --out-dir flye_out
 ↪  --genome-size 2.8g --threads 128
```

#### Polishing

##### Medaka

Medaka was run to polish the assembly.

```
# version 1.11.2
medaka_consensus -i ../p-horse-total-rebasecall.fastq -d
 ↪ assembly.c.fasta.k32.w100.z1000.ntLink.3rounds.fa -o medaka-out -t 32
```

##### Racon

Racon was run but did not result in any improved assembly statistics so this polishing step was reverted in the final assembly.

```
# Run but not used.
racon -t 32 p-horse-total-rebasecall.fastq assembly.ntLink.3rds.racon0.sam.gz
 ↪ assembly.fasta > assembly.racon1.fasta
```

#### Purge_dups

Purge_dups was run to remove haplotigs and contig overlaps in a de novo assembly based on read depth.

```
# Align the data to generate paf files
minimap2 -x map-ont consensus.fasta $i -t 32 | gzip -c - > $i.paf.gz
minimap2 -I 4G -x map-ont -t 32 consensus.fasta
 ↪ p-horse-total-rebasecall.fastq | pigz > p-horse-total-rebasecall.paf.gz

# Produce stats and cutoffs file
./purge_dups/bin/pbcstat p-horse-total-rebasecall.paf.gz
./purge_dups/bin/calcuts PB.stat > cutoffs 2>calcults.log

# Split consensus and self-align
./purge_dups/bin/split_fa consensus.fasta > consensus.split.fa
minimap2 -xasm5 -DP consensus.split.fa consensus.split.fa | gzip -c - >
 ↪ consensus.split.self.paf.gz

# Purge dups and haplotigs
```

```
./purge_dups/bin/purge_dups -2 -T cutoffs -c PB.base.cov
 ↪  consensus.split.self.paf.gz > dups.bed 2> purge_dups.log

# Get purged primary and haplotigs
./purge_dups/bin/get_seqs -e dups.bed consensus.fasta

# Generate histogram, takes a long time. Gives nonsense graph.
./purge_dups/scripts/hist_plot.py -c cutoffs PB.stat PB.base.png
```

#### Curate contigs

Remove mitochondrial contigs, high coverage, and low fragment size contigs

1. Start with `purged.fa` (result of purge_dups)
2. Remove mitogenome: `seqkit grep -v -p "contig_12841_1" purged.fa > output.fasta`
3. Remove low fragment sizes: `seqkit seq -m 10000 output.fasta > output2.fasta`
4. Map reads to assembly: `minimap2 -ax map-ont output2.fasta ../p-horse-total-rebasecall.fastq -t 32 > output2.sam`
5. Sort and index: `samtools sort output2.sam -o output2.bam ; samtools index output2.bam`
6. Remove 15X < all reads > 500X.
    1. `mosdepth -t 32 -n output2 output2.bam`
    2. Look at mosdepth summary and sort by depth
    3. Copy contig names < 500X and > 15X into `keepers.txt`
    4. Sort for uniq contigs: `cat keepers.txt | sort | uniq > keepers.sort.txt`
    5. `seqkit grep -f keepers.sort.txt output2.fasta -o curated.fa`

##### Assembly-stats

```
assembly-stats file.fasta
```

#### Scaffolding (ragtag)

```
# Install ragtag
mamba install -c bioconda ragtag
```

```
# Run it
ragtag.py scaffold ~/Desktop/genomes/GCA_002863925.1_EquCab3.0_genomic.fna
↪ curated.fa -o ragtag-out -t 32
```

#### Scaffolding (ntLink)

ntLink was used to scaffold but did not yield as contiguous results as ragtag so output was not used. It scaffolded and fill gaps in the assembly, reduced contig count, and doubled the N50.

```
# Run ntLink with 3 rounds and gap-filling
ntLink_rounds run_rounds_gaps clean target=assembly.fasta
↪ reads=p-horse-total-rebasecall.fastq k=32 w=100 t=5 rounds=3 overlap=True
```

#### Gap Closing

##### TGS-GapCloser

```
# Run TGS-GapCloser
tgsgapcloser --scaff ragtag.scaffold.fasta --reads
↪ ../p-horse-total-rebasecall.fasta --output ragtag-tgscloser-out --ne
↪ --threads 32
```

#### Visualize scaffolds

##### JupiterPlot

```
# Install circos
sudo apt install circos

# Install JupiterPlot
git clone https://github.com/JustinChu/JupiterPlot

# Create Jupiter plot
```

```
./jupiter name=ragtag
↪ ref=~/Desktop/genomes/GCA_002863925.1_EquCab3.0_genomic.fna
↪ fa=../ragtag.scaffold.fasta

# Use a clean version of the reference horse that only has full chromosomes
seqkit grep -r -p "CM0*" GCA_002863925.1_EquCab3.0_genomic.fna >
↪ GCA_002863925.1_EquCab3.0_genomic.clean.fna

# Visualize contigs (Anything over ng=85 won't render b/c of too many
↪ contigs)
./jupiter name=curated-vs-EquCab3.0
↪ ref=../../GCA_002863925.1_EquCab3.0_genomic.clean.fna
↪ fa=../../curated/curated.fa t=32

# Visualize scaffolds
/jupiter name=tgs-vs-EquCab3.0
↪ ref=../../GCA_002863925.1_EquCab3.0_genomic.clean.fna
↪ fa=../ragtag-tgscloser-out.scaff_seqs t=32 ng=95
```

#### Curate scaffolds

##### Naming

Scaffolds for *Equus ferus przewalskii* (EPR) are assigned chromosome names based on homology to *Equus caballus* (ECA) synteny from EquCab3.0 as shown in Ahrens and Stranzinger 2005. The homologous EPR scaffold to ECA chromosome 5 was split by N gaps into several smaller chromosomes. The largest blocks syntenic to the p- and q-arms were labeled with their chromosome names as shown in the supplementary file.

1. Copy out the ECA syntenic scaffold to a new directory.
2. Split it into a multi-fasta file by all the N gaps. `seqkit seq -w0 ragtag-tgscloser-out.part_CM009152 | awk '{gsub("[Nn]+","\n>\n");}1' | awk '/^>/ {if ($0 == ">") {$0=prev} prev=$0}1' | awk '/^>/ {getline seq} {if(seq!="") {print $0"\n"seq}}' | awk '(/^>/ && a[$0]++) {$0=$0"_"a[$0]}1' > ECAChr5-multi.fa`
3. Split multifasta into separate fasta files. `seqkit split -i ECAChr5-multi.fa`
4. Rename the largest two contigs to either Chr23/24 depending on blast positions and the rest to ChrUn
5. Combine all named Chrs and ChrUns into final scaffold file.

#### Mitochondria

The mitogenome was extracted from the contig assembly.

##### MitoHiFi

1. `sudo docker pull ghcr.io/marcelauliano/mitohifi:master`
2. `singularity shell -bind /home/cfaulk/Desktop/p-horse-analysis/MitoHiFi:/MitoHiFi docker://ghcr.io/marcelauliano/mitohifi:master mitohifi.py -h`
3. `mitohifi.py -c assembly.fasta -f equus_caballus_mtDNA.fa -g equus_caballus_mtDNA.gb -t 32`
4. The "potential contigs" directory lists the contig containing mtDNA (contig_12841).
5. Annotations were saved for submission to NCBI Assembly.

#### Global Methylation

Map the reads and modifications to the assembly and aggregate methylation.

```
# Map
samtools fastq -T "*" p-horse-total-rebasecall.bam | minimap2 -y -L -ax
↪  map-ont curated/EquPr2_scaff.fasta - -t 32 > EquPr2.modmapped.unsort.sam

# Sort and index
samtools sort EquPr2.modmapped.unsort.sam -o EquPr2.modmapped.bam -@ 32 ;
↪  samtools index EquPr2.modmapped.bam -@ 32

# Convert mapped bam file reads to .bed format to quantitate methylation by
↪  position.
~/Desktop/modkit_0.2.2/modkit pileup -t 32 --cpg --ref
↪  ../curated/EquPr2_scaff.fasta --combine-strands --only-tabs
↪  EquPr2.modmapped.bam EquPr2.modkit.bed

# Summarize global methylation
# Report all canonical, 5mc and 5hmc percentages
awk '$4=="m" {can+=$13; mod+=$12; oth+=$14; valid+=$10} END{print (can/valid)
↪  " CpG canonical\n" (mod/valid) " 5mCpG modified\n" (oth/valid) " 5hmCpG
↪  modified"}' EquPr2.modkit.bed
```

#### RepeatMasking

Run RepeatMasker with horse repeats

```
# Run Repeatmasker using Dfan 3.8
/usr/local/RepeatMasker/RepeatMasker -pa 32 -s -xsmall -species horse -dir
↪ repeatmasker-EquPr2 EquPr2_scaff.fasta
```

#### Annotation

Install and run GeMoMa

```
# Install mmseqs2 dependency
mamba install -c conda-forge -c bioconda mmseqs2

# Install GeMoMa
mamba install gemoma -c bioconda

# Prepare to run
#echo "GCF_002863925.1" > horse-GCF.txt
echo "Equus caballus" > horse-GCF.txt
mkdir references
GeMoMa NRR rl=horse-GCF.txt
mv *.gz references

GeMoMa GeMoMaPipeline threads=32 outdir=gemoma-p-horse GeMoMa.Score=ReAlign
↪ AnnotationFinalizer.r=NO o=true t=../curated/EquPr2_scaff.fasta.gz
↪ a=references/GCF_002863925.1_EquCab3.0_genomic.gff.gz
↪ g=references/GCF_002863925.1_EquCab3.0_genomic.fna.gz -Xms5G -Xmx50G

# BUSCO was used to assess completeness in protein mode:
busco -i predicted_proteins.fasta -o busco -m protein -l cetartiodactyla -c
↪ 32
```

#### Variant Calling & Phasing

`Clair3` version 1.0.4 calling `Whatshap` version 1.7 for phasing and `Longphase` version 1.5.2 for haplotagging.

```
# Create environment
mamba create -n clair3 -c bioconda clair3 python=3.9.0 -y
mamba activate clair3

# Download models
git clone https://github.com/nanoporetech/rerio
./rerio/download_model.py --clair3

# Call variants
run_clair3.sh --bam_fn EquPr2.modmapped.bam --ref_fn EquPr2_scaff.fasta
 ↪  --threads 28 --model_path ~/rerio/clair3_models/r1041_e82_400bps_sup_v410
 ↪  --platform ont --output clair3_output
 ↪  --use_whatshap_for_final_output_phasing
 ↪  --use_longphase_for_final_output_haplotagging

# Phasing stats can be seen with whatshap
whatshap stats phased_merged.vcf.gz

# Heterozygosity stats can be determined with the following command
bcftools stats -s - phased_merge_output.vcf.gz

# Transition Transversion ratio can be determined with the following command
vcftools --TsTv-summary --gzvcf phased_merge_output.vcf.gz
```

#### DNA Methylation

##### Extract Methylation Counts

`Modkit` version 0.2.4

```
modkit pileup --log-filepath modkit.log -t 28 -r EquPr2_scaff.fasta --cpg
 ↪  --combine-strands --only-tabs --prefix EquPr2-HP --partition-tag HP
 ↪  phased_output.bam modkit_longphase
```

`modkit` output contains both 5mC and 5hmC. Got 5mC only with `awk -v OFS="\t" '$4 == "m"' HP.bed`.

##### Filtering and Normalization

```r
library(tidyverse) # 2.0.0
library(methylKit) # 1.26.0
library(circlize) # 0.4.15
```

`Methylkit` version 1.26.0

```r
# Specify modkit columns
modkit_cols <- list(
  fraction = FALSE,
  chr.col = 1,
  start.col = 2,
  end.col = 3,
  coverage.col = 5,
  strand.col = 6,
  freqC.col = 11
)

# List modkit files
mk_files <- list.files("~/modkit_out", "5mc-HP[12].bed.gz", full.names = T)
↪  |>
  as.list()

# read in
mk_obj <- methRead(
  mk_files,
  sample.id = mk_files |> map(~ str_extract(.x, "HP[:digit:]")),
  assembly = "EquPr2",
  treatment = c(0, 1),
  pipeline = modkit_cols,
  header = F,
  dbtype = "tabix",
  dbdir = "methylkit_out/tabix",
  mincov = 10
)

# Filter, normalize, unite
mk_meth <- mk_obj |>
  filterByCoverage(lo.count = 10,
    hi.perc = 99.9,
```

```
                 save.db = F) |>
  normalizeCoverage(save.db = F) |>
  unite(save.db = F)
```

##### Call DMRs

Methylation counts tiled 100bp.

```
# DMRs
tile_meth <- tileMethylCounts(mk_meth,
                              win.size = 100,
                              step.size = 100,
                              mc.cores = 8,
                              cov.bases = 10,
                              save.db = F)

tile_diff <- calculateDiffMeth(tile_meth,
                               slim = F,
                               save.db = F)

tile_delta <- getMethylDiff(tile_diff,
                            difference = 50,
                            qvalue = 0.05)
# summary stats
dmr_summary <- getData(tile_delta) |>
  summarize(n_DMR = n(),
    mean_abs_delta = mean(abs(meth.diff)),
    min_delta = min(meth.diff),
    max_delta = max(meth.diff)) |>
  mutate(across(everything(), ~ round(.x, digits = 1))) |>
  bind_cols(data.frame(windows_tested = nrow(getData(tile_diff)),
                       window_size = 100)) |>
  t() |>
  data.frame() %>%
  rename_with(function(x){x = "value"}) |>
  rownames_to_column(var = "stat")
```

##### Closest Genes

Identified closest annotation features to DMRs with `bedtools` version 2.31.1 and `GeMoMa GFFAttributes` subcommand.`bedtools closest -d -t "first" -a dmrs.bed -b gene_attributes.bed > dmr_closest_genes.bed`

```
closest <- read_tsv("dmr_closest_genes.bed",
                    na = c("-1","."),
                    show_col_types = FALSE,
                    col_names = c("chr", "start", "end",
                                  "gene_chr", "gene_start", "gene_end",
                                   "gene",
                                  "dist_bp")) |>
  drop_na(gene) |>
  inner_join(getData(tile_delta), by = c("chr", "start", "end"))
## summary stats
closest |>
  summarize(mean_meth = mean(abs(meth.diff)),
            median_meth = median(abs(meth.diff)))
```

##### Visualize

###### Circlize

`Circlize` package version 0.4.15. Chromosome length data generated with `seqkit fx2tab` version 2.3.0.

Generate data objects for visualization:

```
# chromlengths
chrominfo <- read_tsv("chromInfo_EquPr2.txt",
                      col_names = c("chr", "end"),
                      show_col_types = F) |>
  mutate(start = 1) |>
  relocate(start, .before = end) |>
  filter(str_detect(chr, "Chr") & !str_detect(chr, "Un")) |>
  mutate(chr_num = str_remove(chr, "Chr") |>
         as.numeric()) |>
  arrange(chr_num) |>
  data.frame() # circlize doesn't like tibbles
```

```r
# gene prediction data from GeMoMa GFFAttributes
annot <- read_tsv("gene_attributes.bed",
                  col_names = c("chr", "start", "end", "symbol"),
                  show_col_types = F)
```

```r
# filter for autosomal DMRs near named genes
top_closest <- closest |>
  inner_join(chrominfo, by = "chr") |>
  filter(chr != "ChrX") |>
  arrange(chr_num) |>
  dplyr::select(-chr_num) |>
  filter(!str_detect(gene, "^LOC\\d+")) |>
  filter(dist_bp < 20000)

# multi-dmr windows for nested zooming
spans <- top_closest |>
  group_by(chr, gene) |>
  nest() |>
  mutate(data = map(data, ~ .x |>
                      arrange(start.x))) |>
  mutate(start = map_int(data, ~ pluck(.x, 1, 1) - 500),
         end = map_int(data, ~ pluck(.x, 2, -1) + 500)) |>
  relocate(start:end, .after = chr) |>
  mutate(spans = paste(chr, start, end, sep = "-"),
         width = end - start) |>
  data.frame()

dmr_spans <- spans |>
  dplyr::select(chr:end, spans) |>
  relocate(spans, .before = everything()) |>
  relocate(chr, .after = everything()) |>
  data.frame()

correspondence <- dmr_spans |>
  relocate(c(chr, start, end), .before = everything()) |>
  mutate(start.1 = start,
         end.1 = end) |>
  data.frame()

dmrs <- spans |>
  dplyr::select(-gene, -chr, -start, -end) |>
```

```
  relocate(spans, .before = everything()) |>
  mutate(data = map(data, ~ .x |>
                          dplyr::select(start.x, end.x, meth.diff))) |>
  unnest(data) |>
  data.frame()

# nested gene annotation loci
gene_spans <- spans |>
  dplyr::select(-chr, -start, -end) |>
  relocate(spans, .before = everything()) |>
  mutate(data = map(data, ~ .x |>
                          dplyr::select(start.x, end.x, meth.diff))) |>
  unnest(data) |>
  relocate(gene, .after = end.x) |>
  nest(data = c(-spans, -gene)) |>
  mutate(data = map(data, ~ .x |>
                           arrange(start.x))) |>
  mutate(start = map_int(data, ~ pluck(.x, 1, 1)),
         end = map_int(data, ~ pluck(.x, 2, -1))) |>
  dplyr::select(spans, start, end, gene) |>
  data.frame()

# get single CpG 5mC values per-HP within each dmr_span
by <- join_by(chr, within(x$start, x$end, y$start, y$end))

line_mk <- getData(mk_meth) |>
  inner_join(dmr_spans, by) |>
  mutate(meth1 = numCs1 / (numCs1 + numTs1) * 100,
         meth2 = numCs2 / (numCs2 + numTs2) * 100) |>
  dplyr::select(spans, start.x, end.x, meth1, meth2)

# color objects
set.seed(222)
no_dmr_ind <- which(!chrominfo$chr %in% dmr_spans$chr)
chr_bg_color = rand_color(nrow(chrominfo), transparency = 0.8)
names(chr_bg_color) = chrominfo$chr
chr_bg_color[no_dmr_ind] <- "white"
zoom_colors <- chr_bg_color[correspondence$chr]
col_fun = colorRamp2(breaks = c(0, 50, 100), colors = c("blue", "yellow",
 ↪  "red"))
```

Draw circos plot based on example from Circular Visualizations in R

```r
circos.clear()
# genomic track
f1 = function() {
  circos.par("cell.padding" = c(0, 0, 0, 0),
             start.degree = 90,
             gap.after = c(rep(0, nrow(chrominfo)-2), 2))
  circos.genomicInitialize(
    filter(chrominfo, chr != "ChrX"), plotType = NULL
    )
  circos.track(
    ylim = c(0, 1),
    panel.fun = function(x, y) {
      chr = str_remove(CELL_META$sector.index, "Chr")
      xlim = CELL_META$xlim
      ylim = CELL_META$ylim
      circos.rect(xlim[1], 0, xlim[2], 1)
      circos.text(
        mean(xlim),
        mean(ylim),
        chr,
        cex = 0.9,
        col = "black",
        font = 2,
        facing = "inside",
        niceFacing = TRUE
      )
    },
    track.height = 0.05,
    track.margin = c(0,0),
    bg.border = NULL,
    bg.col = add_transparency(chr_bg_color, 0.5)
  )
}
# nested track
f2 <- function(){
  circos.par(cell.padding = c(0, 0, 0, 0),
             gap.after = c(rep(1, nrow(dmr_spans)))
             )
  circos.genomicInitialize(
    dmr_spans,
    plotType = "axis",
    axis.labels.cex = 0.5 * par("cex"),
```

```
  )
  circos.genomicHeatmap(line_mk, col = col_fun,
                      connection_height = NULL
                      )

  circos.genomicTrack(dmrs,
                    stack = T,
                    panel.fun = function(region, value, ...){
                      circos.genomicRect(region, value, col = "black",
                                         border = NA, ...)
                    },
                    track.height = 0.02,
                    bg.border = NA)

    circos.genomicLabels(gene_spans,
                     labels.column = 4,
                     side = "inside",
                     connection_height = mm_h(2),
                     font = 2
                     )
}
# draw
circos.nested(f1, f2, correspondence,
              connection_col = add_transparency(zoom_colors, 0.5),
              connection_height = mm_h(3), adjust_start_degree = T)
circos.clear()
```

###### MethylArtist

Version 1.2.10

Generate an annotation BED file with the gene symbols for DMR labeling in `methylartist`

Generate gene attribute BED file: `GeMoMa GFFAttributes a=final_annotation.gff f=gene`

Combine with reference gene table from `GeMoMaPipeline` output:

```
ref = read_tsv("reference_gene_table.tabular") %>%
  mutate(genes = str_extract(reference_species_0,
                             "gene-[:alnum:]+") |>
           str_remove("gene-")) %>%
```

```
  dplyr::select(name, genes)

att = read_tsv("GFF_attributes.tabular") %>%
  dplyr::select(chromosome, start = `start position`,
                end = `end postion`, # typo in GeMoMa output cols
                ID)

join = att |>
  inner_join(ref, by = c("ID" = "name")) |>
  dplyr::select(-ID)

write_tsv(join, "gene_attributes.bed",
          col_names = F)
```

Generate 1-column DMR and flank coordinate files for `methylartist` highlight and input regions, respectively:

```
tile_delta_flank <- getData(tile_delta) |>
  mutate(start = if_else(start < 1000,
                         start,
                         start - 1000),
         end = end + 1000) |>
  mutate(interval = paste0(chr, ":", start, "-", end)) |>
  dplyr::select(interval)

tile_delta_intervals <- getData(tile_delta) |>
  dplyr::select(chr, start, end)

head(tile_delta_flank)

write_tsv(tile_delta_flank, col_names = F,
          "tile_delta_flank.txt")
write_tsv(tile_delta_intervals, col_names = F,
          "dmrs.bed")
```

Loop `methylartist locus` to visualize all DMRs, run as `loop_methylartist.sh`:

```
#!/bin/bash

flank=$(cat tile_delta_flank.txt)
```

```
for DMR in $flank
do
methylartist locus \
--ref ../EquPr2_scaff.fasta \
--motif CG \
--mods m \
--primary_only \
--phased \
--ignore_ps \
--color_by_hp \
--phase_labels 1:HP1,2:HP2 \
--statname 5mC \
--samplepalette husl \
--nticks 6 \
--labelgenes \
--highlightpalette "Accent" \
--width 12 --height 10 \
--highlight_bed dmrs.bed
--bed gene_attributes.bed
-b ../clair3_output/phased_output.bam \
--nticks 6 \
-i $DMR
done
```

#### Diploidization

##### BCFtools to make variant diploid fasta file

```
cat EquPr2_scaff.fasta | bcftools consensus ../allin1_clair3_output/phased_merge_output.vcf.g[illegible]
| bgzip > EquPr2_scaff.hap2.fasta.gz
```

### BUSCO

```
# Compleasm as more accurate BUSCO measure
compleasm run -l cetartiodactyla -L ~/Desktop/genomes/mb_downloads -a
↪ ../curated.fa -o compleasm-curated -t 32

# Compleasm in protein mode
```

```
compleasm protein -l cetartiodactyla -L ~/Desktop/genomes/mb_downloads -p
 ↪  predicted_proteins.fasta -o compleasm-protein -t 32

# Run BUSCO as comparison
busco -i curated.fa -o busco_curated -m genome --lineage
 ↪  ~/Desktop/genomes/busco_downloads/lineages/cetartiodactyla -c 32
```

#### Quast

```
# Install
pip install quast

# Run
quast.py --large ../curated/EquPr2_scaff.fasta.gz
 ↪  ../GCA_000696695.1_Burgud_genomic.fna.gz -r
 ↪  ~/Desktop/genomes/GCA_002863925.1_EquCab3.0_genomic.fna --fragmented -o
 ↪  quast-EquPr2 -t 32 --features
 ↪  ../gemoma-out/references/GCF_002863925.1_EquCab3.0_genomic.gff.gz
```
